# PCR-based specific detection of multiple *Nigrospora* species causing a banana leaf spot disease

**DOI:** 10.1101/2025.11.30.691474

**Authors:** Shunsuke Nozawa, Sherman M. Chavez, Aniway F. Penalosa, Kyoko Watanabe

**Author notes:** Corresponding author: (N.S.) and (W.K.).

## Abstract

The genus *Nigrospora* has been identified as the primary causative agent of leaf spot disease in banana plantations in Mindanao, the Philippines. The symptoms caused by *Nigrospora* species are similar to those of black leaf streak disease (black sigatoka), which has made managing the disease challenging. Furthermore, development of rapid and precise molecular diagnostic methodologies for leaf spot disease has remained elusive. In the present study, a polymerase chain reaction (PCR)-based detection system was developed. The system targets a *Nigrospora*-specific genomic region, which was identified through comparative analysis of the genus *Nigrospora* and related genera. The newly designed primer set nigF/nigR specifically detected six pathogenic *Nigrospora* species associated with banana leaf spot disease (*N. chinensis*, *N. lacticolonia*, *N. nigrocolonia*, *N. singularis*, *N. sphaerica*, and *N. vesicularifera*), while no amplicon was observed for 15 non-*Nigrospora* species, including *Pseudocercospora fijiensis* isolated from banana leaves. The results indicate that the primers enable clear differentiation from other genera, including the important pathogen *P. fijiensis*. The primers also detected diverse *Nigrospora* isolates (*N. banbusae*, *N. chinensis*, *N. musae*, *N. oryzae*, *N. sphaerica*, and *N. rubi*) obtained from different hosts and regions within Japan, demonstrating broad applicability. Sensitivity testing demonstrated the capacity to detect as minimal as 10 pg of genomic DNA. Additionally, a pathogenic *Nigrospora* isolate was detected successfully from infected banana tissue. This study presents a novel molecular diagnostic tool for *Nigrospora* species, offering a rapid, specific, and practical approach, which could enhance evidence-based disease management in epidemiological investigations and banana production systems.

## Introduction

The genus *Nigrospora* comprises 15 species currently registered in MycoBank (accessed April 27, 2024) and is regarded as a cosmopolitan fungal group occurring in diverse environments globally (Wang et al. 2017). Conidia of *Nigrospora* spp. are frequently detected in the atmosphere and disperse over long distances through dust storms and other aerial transport events (Wu et al. 2004).

As plant-associated fungi, *Nigrospora* species function both as widespread endophytes (Rashmi 2019; Luo et al. 2017; Safi et al. 2024) and pathogens infecting a broad range of crops, fruit trees, and ornamentals (Wang et al. 2017). Reports of *Nigrospora*-associated diseases have increased globally (Sharma et al. 2013; Liu et al. 2016; Taguiam et al. 2020; Villanueva et al. 2023), highlighting the expanding pathological significance of the genus.

In banana cultivation, *Nigrospora sphaerica* has long been recognized as the causal agent of vascular discoloration and hardening of fruit tissues (McLennan and Hoëttb 1933). In Bangladesh, *N. sphaerica* and *Neocordana musae* were identified as major pathogens responsible for banana leaf spot (Sadid et al. 2023). More recently, however, six *Nigrospora* species (*N. chinensis*, *N. lacticolonia*, *N. nigrocolonia*, *N. singularis*, *N. sphaerica*, and *N. vesicularifera*) have been identified as emerging leaf spot pathogens in banana plantations in Mindanao, Philippines (Nozawa et al. 2025).

Leaf symptoms caused by *Nigrospora* spp. are nearly indistinguishable from those caused by black Sigatoka, one of the most destructive banana diseases globally, which is caused by *Pseudocercospora fijiensis*. The Philippine Banana Industry Roadmap notes the severe impact of major diseases on production (Department of Agriculture 2022), and field observations indicate that *Nigrospora* leaf spot has often been misdiagnosed as black Sigatoka, with fungicides targeting *P. fijiensis* sprayed more than 40 times annually (Department of Agriculture 2022). Our previous in vitro assays revealed that several of the fungicides have low efficacy against *Nigrospora* spp. (Nozawa et al. 2025), suggesting that misdiagnosis could have contributed to shifts in pathogen community structure and facilitated the emergence of *Nigrospora* spp. as dominant pathogens.

Environmental factors can alter foliar fungal communities. Changes in temperature and precipitation have been linked to increases in plant pathogenic guilds (Chen et al. 2021; Dea et al. 2022). Warming can also shift communities toward pathotrophic fungi (Fu et al. 2025; Zhong et al. 2025). Therefore, climate-driven changes in the phyllosphere may also contribute to the emergence of *Nigrospora* leaf spot.

Despite the rising incidence of *Nigrospora*-associated diseases, epidemiological information—including species composition, distribution, and dynamics—remains extremely limited. Whereas specific primers for *Pseudocercospora* spp. are used widely for Sigatoka diagnostics (Arzanlou et al. 2007), no rapid detection method exists for *Nigrospora* spp., representing a major barrier to its accurate diagnosis and management.

Therefore, the objective of the present study was to develop a rapid and specific PCR-based diagnostic method for emerging *Nigrospora* leaf spot pathogens in banana. *Nigrospora*-specific genomic regions were identified through comparative genomics of *Nigrospora* and related genera, new PCR primers were designed, and their specificity, sensitivity, and applicability in field monitoring were evaluated.

## Materials and methods

### Exploring *Nigrospora*-specific genes and primer design

Genome assemblies of 14 strains of *Nigrospora* (13 species), 13 species of *Apisospora*, and two species of *Arthrinium*, *Neoarthrinium moseri*, and *Seiridium cupressi* were obtained from the NCBI database (https://www.ncbi.nlm.nih.gov/; Table 1). Gene prediction was conducted using Augustus v3.3.3 (https://github.com/Gaius-Augustus/Augustus) with the *Fusarium* gene model. Single-copy orthologous genes were identified through reciprocal BLASTP searches (BLAST 2.9.0+; https://blast.ncbi.nlm.nih.gov/Blast.cgi; E-value ≤1e–5). Genes present only in *Nigrospora* spp. were screened by comparison with non-*Nigrospora* genomes, and candidate regions were checked further by BLAST searches against the NCBI database. To verify that *Nigrospora* strains possessing the primer-targeted genomic region formed a monophyletic group, a genome-scale phylogenetic analysis was conducted (Supplementary Figure 1). Briefly, orthologous genes were aligned, trimmed, concatenated, and analyzed using the neighbor-joining method with *S. cupressi* as the outgroup using nucleotide data of 2,089 genes (3,688,761 bp).

**Table 1.** List of GenBank accession numbers of genome data.

| Species | Strain | GenBank accession no. |
| --- | --- | --- |
| *Apiospora arundinis* | AAU 773 | GCA_039105305 |
| *Ap. Aurea* | CBS 24483 | GCA_038362705 |
| *Ap. guizhouensis* | F035 | GCA_040333125 |
| *Ap. hydei* | CBS 114990 | GCA_038363865 |
| *Ap. kogelbergensis* | CBS 113332 | GCA_038363985 |
| *Ap. koreana* | KUC21332 | GCA_017163955 |
| *Ap. malaysiana* | STlab-iicb | GCA_006508115 |
| *Ap. marii* | CBS 49790 | GCA_038362585 |
| *Ap. phragmitis* | CBS 135458 | GCA_038363925 |
| *Ap. pterosperma* | CBS 123185 | GCA_019976485 |
| *Ap. rasikravindrae* | CBS 33761 | GCA_038363885 |
| *Ap. saccharicola* | CBS 83171 | GCA_038363915 |
| *Ap. sphaerosperma* | JCM 6018 | GCA_040366785 |
| *Arthrinium phaeospermum* | M.B.Ellis | GCA_006503535 |
| *Ar. puccinioides* | CBS 549.89 | GCA_022414665 |
| *Neoarthrinium moseri* | TUCIM 5799 | GCA_022829195 |
| *Nigrospora bambusae* | LC7114 | GCA_048773625 |
| *N. camelliae-sinensis* | LC3500 | GCA_048773585 |
| *N. chinensis* | LC4575 | GCA_048773525 |
| *N. guilinensis* | LC3481 | GCA_048773485 |
| *N. hainanensis* | LC7030 | GCA_048773445 |
| *N. laticolonia* | LC3324 | GCA_048773465 |
| *N. musae* | Guizhou | GCA_048773425 |
| *N. oryzae* | GZL1 | GCA_016758845 |
| *N. osmanthi* | Z1 | GCA_029489945 |
| *N. pyriformis* | LC2045 | GCA_048773265 |
| *N. rubi* | LC2698 | GCA_048773245 |
| *N. sphaerica* | LC2840 | GCA_048773285 |
|  | ZJJ_C1 | GCA_018287875 |
| *N. vesicularis* | LC7010 | GCA_048773225 |
| *Seiridium cupressi* | BM-138-000234 | GCA_036674765 |

One conserved *Nigrospora*-specific gene was selected based on its presence across genomes of *Nigrospora* species and absence from all other genera. Primers nigF (5’-TCG GGC CTC TGG CGG ACC GA-3’) and nigR (5’ -ATG CCR AAG CTC GTR ATG AA-3’) were designed, producing a 425–428-bp amplicon (Fig. 1).

**Fig. 1.**
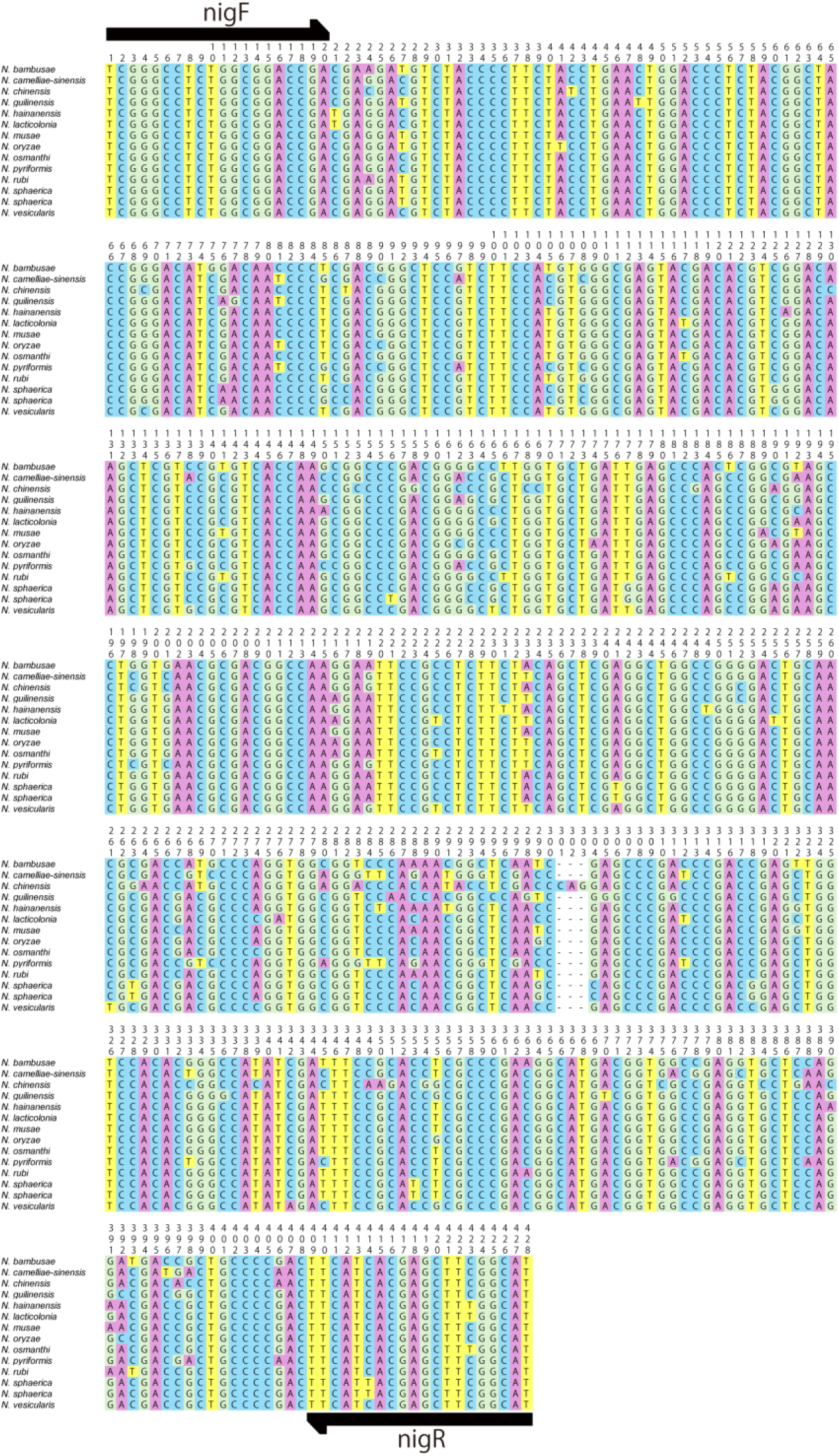
Primer binding sites of nigF and nigR. Highlighted regions indicate the binding sites of primers nigF and nigR on the target gene sequence.

### Evaluation of primer specificity

To assess primer specificity, six pathogenic *Nigrospora* species associated with banana leaf spot (*N. chinensis, N. lacticolonia, N. nigrocolonia, N. singularis, N. sphaerica,* and *N. vesicularifera*), six *Nigrospora* species isolated in Japan (*N. bambusae, N. chinensis, N. musae, N. oryzae, N. rubi,* and *N. sphaerica*), 57 fungal strains belonging to 14 genera collected from banana leaves on Mindanao Island, Philippines (Table 2), and *P. fijiensis* from a diseased banana leaf isolated in Japan were tested. Mycelia were grown on Potato Dextrose Agar (PDA) for 3–20 days, and genomic DNA was extracted using the CTAB method (Doyle and Doyle 1987). PCR was performed in 10-μL reactions containing 1 μL DNA, 1 μL 10× Ex Taq buffer (with MgCl₂), 0.8 μL dNTPs (10 mM each), 0.1 μL each primer (50 μM), 0.05 μL Ex Taq polymerase (Takara), and 7 μL sterile water. Amplification of the Internal Transcribed Spacer (ITS) region using ITS5/ITS4 (White et al. 1990) confirmed success of DNA extraction. The PCR cycle for nigF/nigR consisted of 94°C for 5 min, 35 cycles of 94°C for 30 s, 69°C for 20 s, 72°C for 30 s, and 72°C for 3 min. PCR products were purified using ExoSAP-IT reagent (Thermo Fisher Scientific, Waltham, MA, USA). Sequencing was conducted using the primer nigF by Fasmac Inc. (Atsugi, Japan). The obtained sequences were compared with the target region of *N. lacticolonia* (Strain: LC3324; GenBank accession no.: GCA_048773465) to confirm that the amplified sequences were the target sequence. The PCR test was examined twice.

**Table 2.** Strains used for PCR tests.

| Genus | Isolates | Source | Origin country |
| --- | --- | --- | --- |
| *Alternaria* sp. | TAP24N072 | banana | Philippines |
| *Colletotrichum* sp. | TAP24N009 | banana | Philippines |
| *Colletotrichum* sp. | TAP24N020 | banana | Philippines |
| *Colletotrichum* sp. | TAP24N032 | banana | Philippines |
| *Colletotrichum* sp. | TAP24N061 | banana | Philippines |
| *Colletotrichum* sp. | TAP24N080 | banana | Philippines |
| *Curvularia* sp. | TAP24N052 | banana | Philippines |
| *Curvularia* sp. | TAP24N152 | banana | Philippines |
| *Dimorphiseta* sp. | TAP24N170 | banana | Philippines |
| *Exserohilum* sp. | TAP24N130 | banana | Philippines |
| *Fusarium* sp. | TAP24N015 | banana | Philippines |
| *Fusarium* sp. | TAP24N042 | banana | Philippines |
| *Fusarium* sp. | TAP24N044 | banana | Philippines |
| *Fusarium* sp. | TAP24N046 | banana | Philippines |
| *Fusarium* sp. | TAP24N084 | banana | Philippines |
| *Fusarium* sp. | TAP24N116 | banana | Philippines |
| *Fusarium* sp. | TAP24N117 | banana | Philippines |
| *Fusarium* sp. | TAP24N133 | banana | Philippines |
| *Fusarium* sp. | TAP24N135 | banana | Philippines |
| *Fusarium* sp. | TAP24N144 | banana | Philippines |
| *Fusarium* sp. | TAP24N147 | banana | Philippines |
| *Fusarium* sp. | TAP24N173 | banana | Philippines |
| *Lasiodiplodia* sp. | TAP24N064 | banana | Philippines |
| *Microdochium* sp. | TAP24N114 | banana | Philippines |
| *Microdochium* sp. | TAP24N142 | banana | Philippines |
| *Microdochium* sp. | TAP24N148 | banana | Philippines |
| *Microdochium* sp. | TAP24N175 | banana | Philippines |
| *Musicillium* sp. | TAP24N025 | banana | Philippines |
| *Mycosphaerella* sp. | TAP24N121 | banana | Philippines |
| *Neosetophoma* sp. | TAP24N097 | banana | Philippines |
| *Neosetophoma* sp. | TAP24N111 | banana | Philippines |
| *Neosetophoma* sp. | TAP24N140 | banana | Philippines |
| *Neosetophoma* sp. | TAP24N155 | banana | Philippines |
| *Neosetophoma* sp. | TAP24N159 | banana | Philippines |
| *Neosetophoma* sp. | TAP24N172 | banana | Philippines |
| *Neosetophoma* sp. | TAP24N178 | banana | Philippines |
| *Nigrospora bambusae* | MAFF 235566 | wheat | Japan |
| *N. chinensis* | TAP24N049 | banana | Philippines |
| *N. chinensis* | MAFF 425112 | oshimazakura | Japan |
| *N. lacticolonina* | TAP24N001 | banana | Philippines |
| *N. musae* | MAFF 425574 | japanese gentian | Japan |
| *N. nigrocolonina* | TAP24N202 | banana | Philippines |
| *N. singularis* | TAP24N034 | banana | Philippines |
| *N. oryzae* | MAFF 111239 | rice | Japan |
| *N. rubi* | MAFF 425114 | hinoki cypress | Japan |
| *N. sphaerica* | TAP24N107 | banana | Philippines |
| *N. sphaerica* | MAFF 425575 | japanese gentian | Japan |
| *N. vesicularifera* | TAP24N023 | banana | Philippines |
| *Ochrocladosporium* sp. | TAP24N106 | banana | Philippines |
| *Phaeosphaeria* sp. | TAP24N007 | banana | Philippines |
| *Phaeosphaeria* sp. | TAP24N037 | banana | Philippines |
| *Phaeosphaeria* sp. | TAP24N059 | banana | Philippines |
| *Phaeosphaeria* sp. | TAP24N066 | banana | Philippines |
| *Phaeosphaeria* sp. | TAP24N075 | banana | Philippines |
| *Phaeosphaeria* sp. | TAP24N083 | banana | Philippines |
| *Phaeosphaeria* sp. | TAP24N087 | banana | Philippines |
| *Phaeosphaeria* sp. | TAP24N089 | banana | Philippines |
| *Phaeosphaeria* sp. | TAP24N109 | banana | Philippines |
| *Phaeosphaeria* sp. | TAP24N124 | banana | Philippines |
| *Phaeosphaeria* sp. | TAP24N138 | banana | Philippines |
| *Phaeosphaeria* sp. | TAP24N151 | banana | Philippines |
| *Phaeosphaeria* sp. | TAP24N153 | banana | Philippines |
| *Phaeosphaeria* sp. | TAP24N160 | banana | Philippines |
| *Phaeosphaeria* sp. | TAP24N162 | banana | Philippines |
| *Phaeosphaeria* sp. | TAP24N169 | banana | Philippines |
| *Pseudopestalotiopsis* sp. | TAP24N051 | banana | Philippines |
| *Pseudopestalotiopsis* sp. | TAP24N119 | banana | Philippines |
| *Pseudocercospora fijiensis* | MAFF 237907 | banana | Japan |
| *Zasmidium* sp. | TAP24N004 | banana | Philippines |

### Primer sensitivity

Sensitivity of the primer set was evaluated using serially diluted genomic DNA of *N. lacticolonia* (TAP24N011) at concentrations of 0.1 pg/μL, 1 pg/μL, 10 pg/μL, 100 pg/μL, 1 ng/μL, and 10 ng/μL. PCR was performed under the conditions described for specificity testing. The lowest concentration yielding a visible band was recorded as the detection limit. The PCR test was examined twice.

### Detection of *Nigrospora* in banana leaf tissues

Banana leaves (cv. Dwarf Cavendish) were wounded and inoculated with 5-mm PDA plugs of *N. lacticolonia* (TAP24N011). Inoculated leaves were maintained under high humidity at approximately 25°C for 3 d. Lesion tissues (0.2 g) and healthy controls were surface-sterilized (70% ethanol ×2, sterile water ×2) and subjected to DNA extraction using the CTAB method. DNA quality was confirmed by amplification of the banana actin gene using primers musa_act1F (5’-TGG TGT CAT GGT TGG GAT GG-3’) and musa_act1R (5’-ATT TCC CGT TCA GCA GTG GT-3’) (575 bp). Detection of *Nigrospora* was performed using nigF/nigR under the PCR conditions described above. DNA from PDA-inoculated sites and sterile water served as negative controls; DNA from *N. lacticolonia* mycelia served as a positive control. Amplicons were resolved on 1.4% agarose gels.

## Results

### Primer specificity and sensitivity

Our PCR testing showed that our primers nigF and nigR amplified approximately 400 bp of *Nigrospora* spp. from banana leaves from the Philippines and another host from Japan, but not other strains belonging to other 57 strains of 15 genera isolated from banana leaves obtained in the Philippines and *P. fijiensis*, which is another key pathogen of banana leaf, isolated in Japan (Table 2; Fig. 2). Success of DNA extraction for all strains was confirmed based on the PCR result for ITS regions. Sequence analyses showed sequences from *Nigrospora* spp. were matched with target sequences retrieved from genome data of *N. lacticolonia,* with 85.7 to 99.17% identity (Supplementary Table 1).

**Fig. 2.**
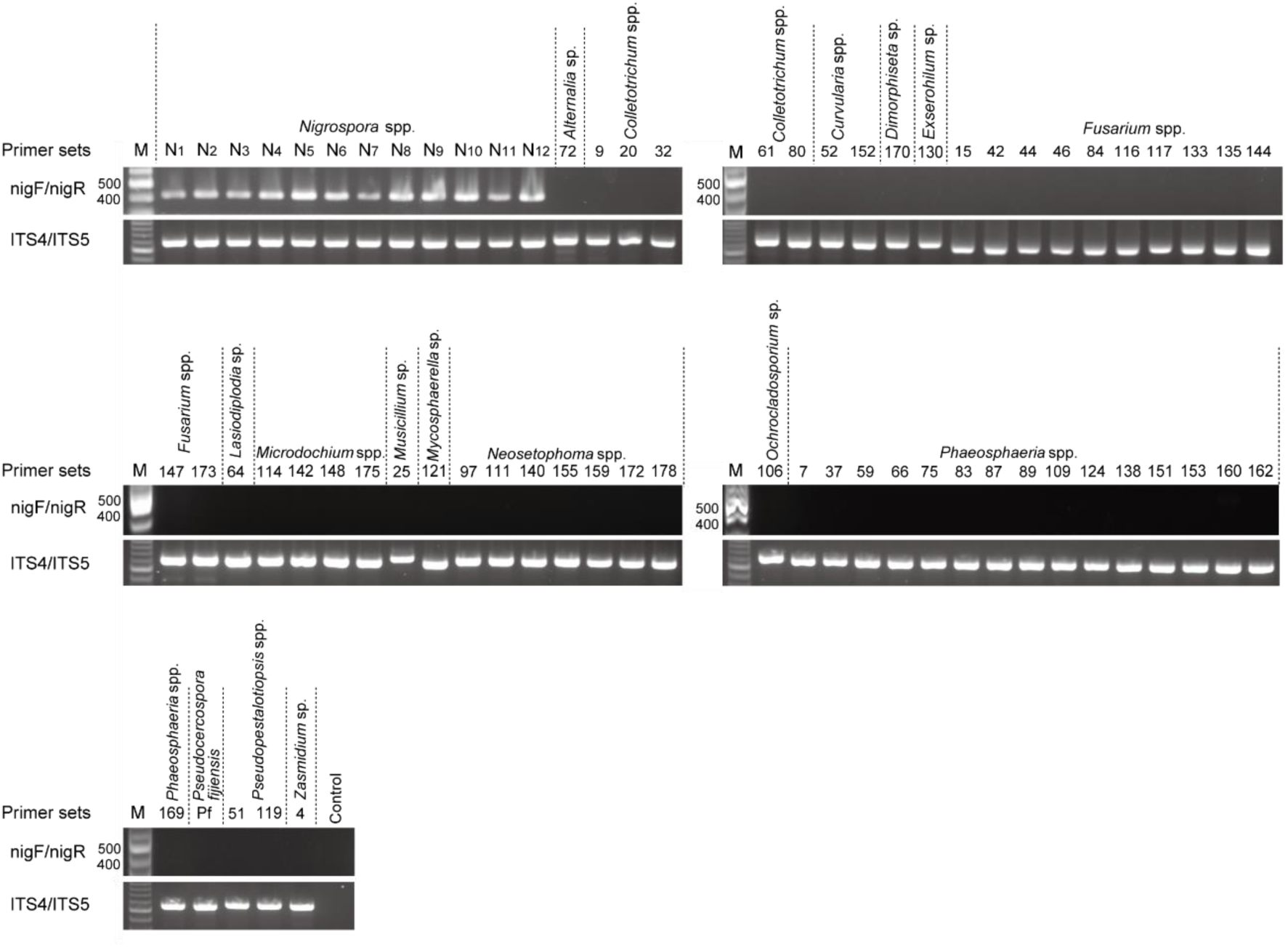
PCR amplification of *Nigrospora* species and other fungal genera isolated from banana leaves using the nigF/nigR primer set. Species names corresponding to each lane are indicated above the gels. Lane M, 100-bp DNA ladder. The upper panel shows amplification of the *Nigrospora*-specific target region, and the lower panel shows amplification of the ITS region as a control. Lanes labeled “N” correspond to *Nigrospora* strains: N1, *N. banbusae* (MAFF 235566); N2, *N. chinensis* (TAP24N049); N3, *N. chinensis* (MAFF 425112); N4, *N. lacticolonia* (TAP24N001); N5, *N. musae* (MAFF 425574); N6, *N. nigrocolonia* (TAP24N202); N7, *N. oryzae* (MAFF 111239); N8, *N. singularis* (TAP24N034); N9, *N. sphaerica* (TAP24N107); N10, *N. sphaerica* (MAFF 425575); N11, *N. rubi* (MAFF 425114); and N12, *N. vesicularifera* (TAP24N023). Numbers above lanes indicate abbreviated strain codes beginning with “TAP24N,” as listed in Table 2. Lane NTC, no-template control (ddH₂O).

The detection limit of the developed primers was examined using serially diluted *N. lacticolonia* (TAP24N011) DNA (Fig. 3). The primer set amplified target size of DNA fragments at concentrations of 10 pg/µl, 100 pg/µl, 1 ng/µl, and 10 ng/µl, which correspond to 0.2 to 2×10^2^ genome copies of *N. lacticolonia*, estimated based on genome assembly data (Genbank accession no. GCA_048773465; genome size: 45.8 Mb). No amplicons were observed at 1 pg/µl, 0.1 pg/µl, and in the negative control.

**Fig. 3.**
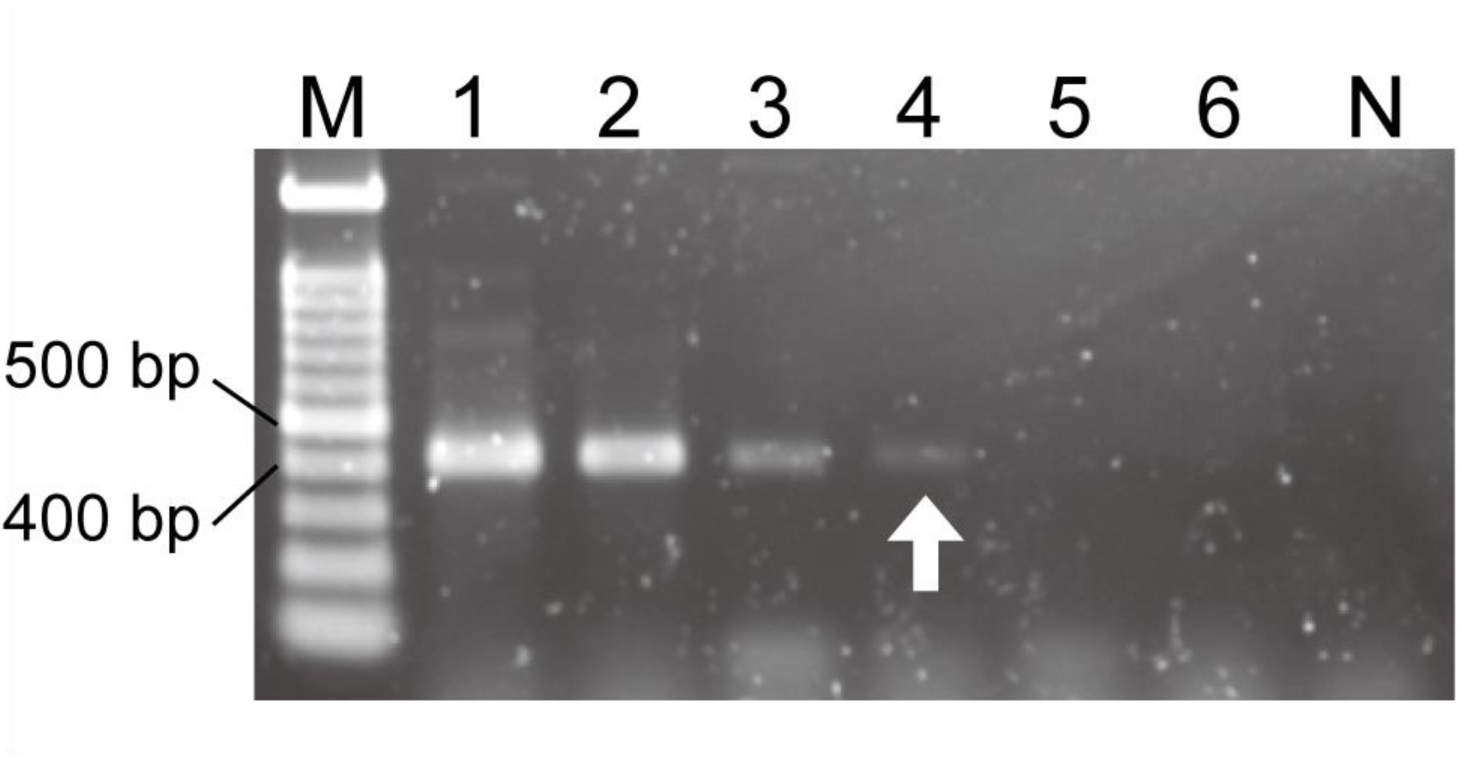
Sensitivity of the *Nigrospora*-specific primer set nigF/nigR. Genomic DNA of *Nigrospora lacticolonia* (TAP24N011) was serially diluted from 10 ng to 0.1 pg and subjected to PCR amplification. Lane M, 100-bp DNA ladder; lanes 1–6, PCR amplicons obtained from 10 ng, 1 ng, 100 pg, 10 pg, 1 pg, and 0.1 pg of template DNA, respectively; lane N, no-template control (NTC). The arrow indicates the lowest detectable DNA concentration.

### Detection from tissues

To verify whether the primer pair nigF and nigR can detect the pathogen infecting banana leaf tissues, PCR was performed using DNA templates derived from banana leaf lesions artificially infected with *N. lacticolonia* (TAP24N011) and from banana leaf tissue inoculated with PDA medium alone (Fig. 4a, b). The approximately 400-bp PCR amplicons were only obtained from DNA extractions from diseased tissues experimentally infected by *N. lacticolonia* (n = 4) and mycelia of *N. lacticolonia* (Fig. 4c). No PCR amplicons were obtained from healthy tissues (n = 4) or sterile deionized water. The success of DNA extraction from plant tissues was verified through PCR targeting the *ACT1* gene region of banana (Fig. 4d).

**Fig. 4.**
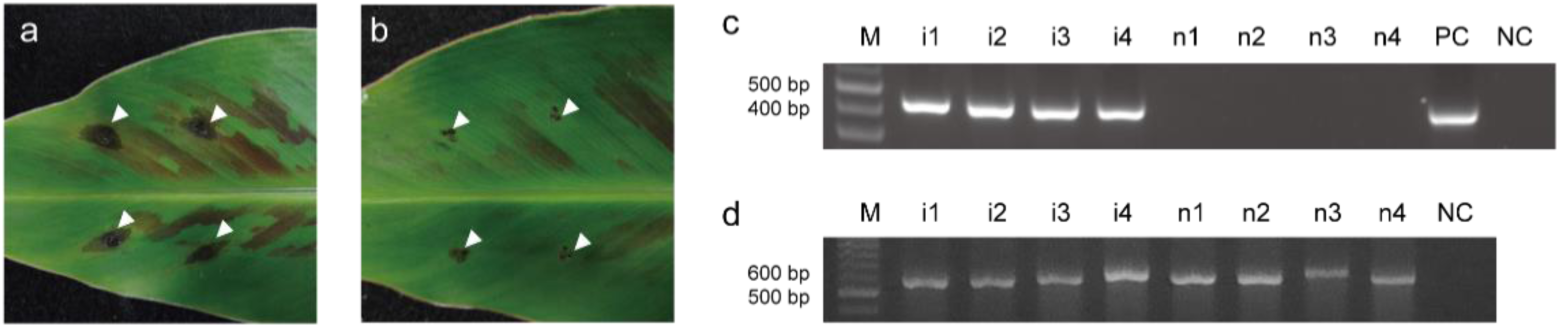
PCR-based detection of *Nigrospora lacticolonia* from banana leaf tissues. (a) Inoculated and (b) non-inoculated leaves of *Musa acuminata* ‘Dwarf Cavendish’. White arrows indicate sites where mycelial or Potato Dextrose Agar disks were placed, and from which template DNA was extracted. (c) PCR amplification using the *Nigrospora*-specific primer pair nigF/nigR from diseased tissues, healthy tissues, and mycelia of *Nigrospora lacticolonia* (TAP24N011). (d) PCR amplification of the ACT1 gene as a control from diseased and healthy tissues. Lane M, 100-bp DNA ladder; lanes i1–i4, amplicons from four inoculated sites; lanes n1–n4, amplicons from four non-inoculated sites; lane NTC, no-template control; lane PPC, positive control from fungal mycelia.

## Discussion

In the present study, a rapid and specific PCR-based detection method for *Nigrospora* species, the causative agent of emerging leaf spot outbreaks in banana plantations on Mindanao Island, the Philippines, was developed. Although specific primers for *P. fijiensis*, the causal agent of black Sigatoka, have been adopted widely for routine monitoring (Arzanlou et al. 2007), no comparable diagnostic tool has been available for *Nigrospora* spp. (Nozawa et al. 2025). The similarity between leaf symptoms caused by *Nigrospora* spp. and those caused by black Sigatoka has likely contributed to management failures in affected plantations, with misdiagnosis and inappropriate fungicide application as potential consequences.

The six pathogenic *Nigrospora* species isolated from Mindanao Island establish a phylogenetic assemblage consistent with previous taxonomic frameworks (Wang et al. 2017; Manawasinghe et al. 2024; Nozawa et al. 2025), comprising *N. chinensis* at the basal position and five sister species. In light of the observed broad phylogenetic diversity, a set of primers was designed to target a specific genomic region of the genus *Nigrospora* that is absent in the closely related genus *Apisospor*a. The newly developed nigF/nigR primers successfully amplified all six pathogenic species associated with banana leaf spot (*N. chinensis*, *N. lacticolonia*, *N. nigrocolonia*, *N. singularis*, *N. sphaerica*, and *N. vesicularifera*), while no amplification occurred in 15 non-*Nigrospora* genera isolated from banana leaves, demonstrating high specificity.

Furthermore, the primers demonstrated the capacity to amplify *Nigrospora* isolates derived from various host species in Japan, signifying their broad applicability across diverse hosts and geographic origins. Notably, the primers did not amplify the primary banana pathogen, *P. fijiensis*, enabling reliable differentiation in fields where multiple pathogens may co-occur.

The sensitivity assay demonstrated that the nigF/nigR primer set can detect as little as 10 pg of genomic DNA (approximately 0.2 genome copies). Furthermore, infected banana tissues yielded positive amplification without the need for pathogen isolation. The finding signifies a substantial practical advantage, as *Nigrospora* spp. have traditionally posed a significant challenge with regard to identification, attributable to the limited species resolution of the ITS region (Wang et al. 2017). The present study offers the first practical molecular marker capable of overcoming such diagnostic limitations.

From a disease-management perspective, the availability of a rapid detection tool for *Nigrospora* spp. has significant implications. Misdiagnosis of *Nigrospora* leaf spot as black Sigatoka can result in excessive and ineffective fungicide applications, while concomitantly increasing fungicide resistance risk in *P. fijiensis*. By facilitating on-site differentiation between the two diseases, the primer set could optimize fungicide utilization, minimize superfluous applications, and bolster evidence-based management decisions in banana plantations.

*Nigrospor*a species are also significant pathogens of various other crops, including *Brassica juncea* (Sharma et al., 2013), tea (Liu et al., 2016), dragon fruit (Taguiam et al., 2020), and cacao (Villanueva et al., 2023). *N. oryzae* is a causative agent of grain discoloration and panicle blight in rice (Savino and Caretta 1992), and it has been misdiagnosed as rice blast, resulting in the implementation of inappropriate control measures (Liu et al. 2021).

In light of such findings, the nigF/nigR primers developed in the present study could also be applicable to *Nigrospora* diseases in other crops, pending appropriate validation. The present study is not without limitations. Primer validation was conducted primarily using isolates from the Philippines and Japan; further testing is necessary to confirm applicability to populations from other regions. Secondly, the efficiency of PCR amplification can be influenced by the thermal characteristics of various PCR platforms, as the heating and cooling behavior of different thermal cyclers can vary. Considering the validation was performed using a single cycler model, further assessment across multiple PCR instruments is necessary before large-scale deployment of the diagnostic tool.

## Data Availability

The datasets generated during the current study are available in Supplementary data.

## Author Contribution

S.N. conceived the study, designed the methodology, conducted all analyses, visualized the data, and led manuscript preparation. K.W. supervised the study, acquired funding, and assisted in manuscript editing and project coordination. All authors have reviewed and approved the final manuscript.

