## Supplementary materials for "PCR-based specific detection of multiple *Nigrospora* species causing a banana leaf spot disease"

**Supplementary Material**

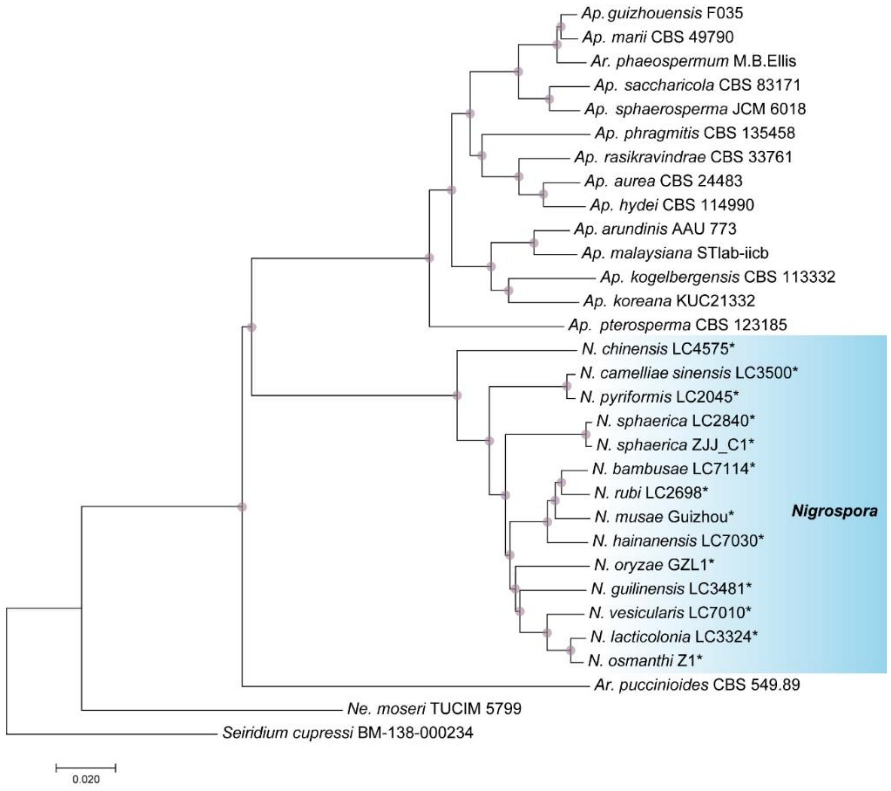


**Supplementary Figure 1. Genome-scale phylogeny of *Nigrospora* and related genera.** Neighbor-joining tree based on concatenated nucleotide alignments of 2,089 single-copy orthologous genes. Purple circles on branches indicate 100% bootstrap support. The *Nigrospora* clade is highlighted in blue. Strains marked with asterisks possess the gene region targeted by primers nigF and nigR. *Seiridium cupressi* was used as the outgroup.

**Supplementary Table 1** Identities between target sequences retrieved from genome data of *N. lacticolonia* and sequences of PCR products.

| Species (strain) | Identity (%) |
|---|---|
| *N. banbusae* (MAFF 235566) | 93.7 |
| *N. chinensis* (MAFF 425112) | 85.7 |
| *N. chinensis* (TAP24N249) | 85.7 |
| *N. lacticolonia* (TAP24N011) | 99.17 |
| *N. musae* (MAFF 425574) | 95.17 |
| *N. nigrocolonia* (TAP24N202) | 95.4 |
| *N. oryzae* (MAFF 111239) | 95.7 |
| *N. rubi* (MAFF 425114) | 94.57 |
| *N. singularis* (TAP24N034) | 88.2 |
| *N. sphaerica* (MAFF 425575) | 93.1 |
| *N. sphaerica* (TAP24N107) | 93.1 |
| *N. vesicularis* (TAP24N023) | 93.4 |
